# Repeated stress persistently suppresses the central node in the courtship circuitry in *Drosophila*

**DOI:** 10.64898/2025.12.14.693558

**Authors:** Yuto Nishizuka, Masato Tsuji, Kazuo Emoto

## Abstract

Stress caused by external stressors, such as painful stimulation, can elicit a drastic set of behavioral responses. Such stress responses normally decay soon after the stressor disappears, but can become persistent when the stressor becomes longer and/or repeated. The neuronal mechanism underlying the persistence of stress responses, however, remains elusive. Here, we discovered that repeated electric shocks persistently suppress male Drosophila’s courtship behavior. We comprehensively evaluated the physiological changes in the genetically defined courtship circuitry, and found that a courtship-triggering node of the circuitry, called P1 neurons, is preferentially and persistently suppressed following electric shocks. Our data uncovers in a comprehensively defined courtship circuitry that repeated stress persistently suppresses a critical node of the circuitry in a targeted fashion to alter behavior.

## INTRODUCTION

Stress caused by external stressors, such as painful stimulation, can elicit a drastic set of behavioral responses accompanied by neuronal and molecular signatures in the brain^123^. Such stress responses normally decay soon after the stressor disappears, but can become chronic when the stressor becomes longer and/or more repeated^1–3^. Previous literature, mostly using rodent as a model, has uncovered the correlates of such chronic stress responses in the brain. For instance, repeated exposure to electric shocks modulates neuronal activities, synaptic plasticity, transcriptional and epigenetic regulations^3–8^. All these modulations persist after the shocks terminate for days or weeks, just like the behavioral changes ^3–8^. Reversing / artificially inducing some of these modulations can reduce / mimic stress-induced behavioral changes^3,5,8^. Thus, neuronal mechanism underlying long-term/repeated stress-induced behavioral changes have seemingly been partially resolved. However, since in virtually all cases the neuronal circuitry of relevant behaviors is only partially known, we do not yet know whether additional modulations take place in the relevant circuitry outside the neurons that have so far been tested, nor do we know how such modulation might impact the physiology of the overall circuitry as a network.

Drosophila male courtship circuitry is probably one of the best well-studied neuronal circuitries, in which the information flow from the first order sensory neurons to the motor neurons has been structurally and physiologically traced^9,10^. In this circuitry, the first order gustatory neurons in the foreleg detect the presence of female-specific pheromone 7,11-HD^11^. These neurons then project to the second order neurons in the brain, called vAB3 and mAL^9^. vAB3 activates a group of third order neurons called P1 neurons^9^. mAL, in the meantime, provides inhibitory input to P1, thus implementing a feed-forward inhibition^9,12^. Activation of P1 neurons triggers the whole, complex set of courtship behaviors, such as orienting, singing a love song by extending and vibrating the wing, and mounting^10,13^. P1 triggers singing through its projection to pIP10 neurons, which then project to the ventral nerve cord^14^. In the ventral nerve cord, a group of neurons called vPR6 receive this input and project to a set of motor neurons to drive singing^14^. Thus, we reasoned that, if repeated stress suppresses courtship, as has been reported in mammals ^15–18^, it would be feasible to reveal the mechanism underlying behavioral changes in the context of the whole courtship circuitry.

Indeed, Drosophila exhibits behavioral changes due to repeated stress. For example, repeated vibration stress can reduce courtship and “gap-climbing” behaviors^19^. At least the gap-climbing reduction can be rescued by feeding the flies with high-concentration sugar^20^, or clinically prescribed anti-depressants, serotonin-specific reuptake inhibitor or lithium chloride^19^. Such stress/sugar effects have been mapped to a specific subtype of neurons innervating the mushroom body and a serotonergic signaling there^19^. Repeated rejection by a non-virgin female over a few days can also suppress courtship behavior^21^. Similar phenomenon has been reported for repeated losing to a hyper-aggressive male in fights^22^. However, as in mammalian studies, we currently lack understanding of how repeated stress results in behavioral changes in the context of the whole of the relevant neuronal circuitry.

In the present study, we set out to test how repeated stress might modulate the courtship circuitry, and sought to understand what drives such a modulation to alter behavior.

## RESULTS

### **I.** Repeated electric shocks suppress courtship

We first sought to test if repeated stress suppresses courtship. To this end, we developed an electric shock chamber, in which an array of copper wires tiles the floor, so that “+” and “-“ wires alternate each other (Fig. 1a; Supplementary Fig. 1a). First, a group of 10 flies are transferred into the chamber. Soon after, a 3-hour shock protocol has been initiated, in which the copper wires are now charged, but the current flows only when a fly steps over both “+” and “-“ wires. This resulted in sparse shocks at random timings, inducing flying, jumping, and running behaviors (Supplementary Fig. 1b; Supplementary Movie 1). This set of behaviors is typically observed in response to a visual loom or a heat shock of > 42^23–25^, consistent with these behaviors being escape attempts. After approximately 10 min, however, the frequency of shocks sharply declined, and the flies tended to stand still, avoiding to step over “+” and “-“ wires. After 3 hours, the flies were transferred back to the food containing vials and were allowed to recover until the shocks resumed the next day. We implemented the shock protocol for 1, 2, 3, 4 days, and tested the courtship behavior 1 day afterward (Fig. 1b).

**Fig. 1.**
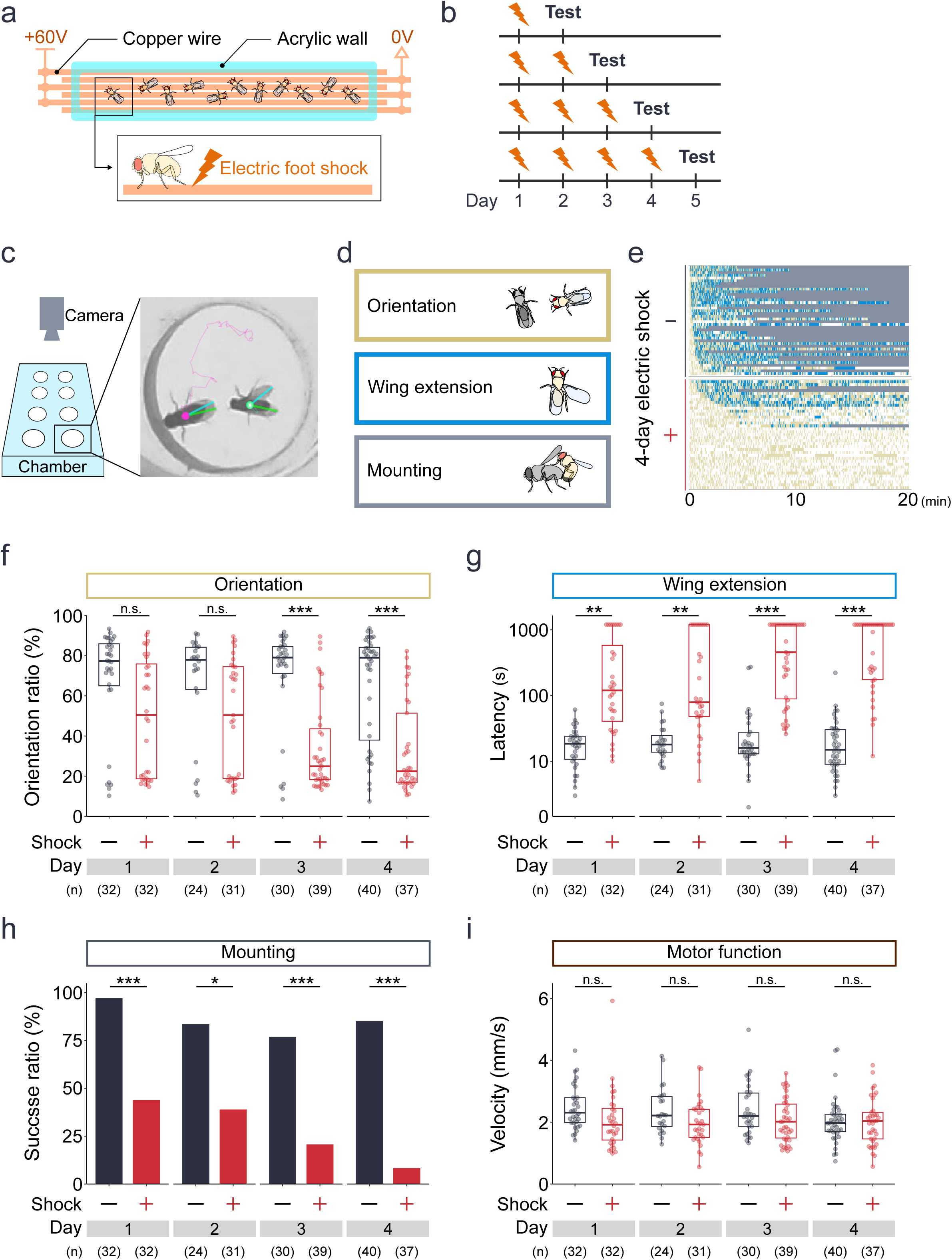
Repeated electric shocks suppress courtship. **a.** An electric shock chamber. An array of copper wires tiles the floor, so that “+” and “-“ wires alternate each other. The copper wires are charged, but the current flows only when a fly steps over both “+” and “-“ wires. **b.** Time course of the experiment. Flies were exposed to electric shocks for 3 hours per day. We implemented the shock protocol for 1, 2, 3, 4 days, and tested the courtship behavior 1 day afterward. **c.** Courtship assay. A male was paired with a female conspecific in a small chamber, and his behavior was continuously recorded for 20 min. **d.** Courtship behaviors quantified in the present study. **e.** Ethogram of color-coded courtship behaviors. Gold, blue, gray indicates orienting, singing, and mounting, respectively. When orienting and singing were detected in the same frame, only singing is shown in this plot. **f.** % time spent on orientation between the beginning of the recording and the onset of mounting (or the end of recording, if the male failed to mount). ***p < 0.001, n.s.: p > 0.05, two-way ANOVA followed by Tukey HSD. **g.** Latency of the initial wing extension in seconds. ***p < 0.001, **p < 0.01, two-way ANOVA followed by Tukey HSD. **h.** % flies mounted a female. ***p < 0.001, *p < 0.05, two-way ANOVA followed by Tukey HSD. **i.** The average locomotion velocity, in mm/s, from the beginning of the recording to the onset of mounting. n.s.: p > 0.05, two-way ANOVA followed by Tukey HSD.

We assessed the male’s courtship behavior following the shock conditioning. Specifically, each male was paired with a female conspecific in a small chamber, and his behavior was continuously recorded for 20 min (Fig. 1c; Supplementary Fig. 1c). We discovered that, while the control flies exhibited bouts of orienting, singing (“wing extension”), and mounting vigorously, the conditioned flies failed to show similar vigor, both in terms of the amount of time spent on, and the latency of initiating, each behavior (Fig. 1d, e; Supplementary Movie 2). We also found that the longer the conditioning, the more pronounced the reduction in courtship became (Fig. 1f-h). The reduction in courtship behavior was likely not due to disrupted motor capabilities inflicted by electric shocks, as the walking speed during courtship behavior did not significantly differ between control and conditioned flies (Fig. 1i). Our data suggests that repeated electric shocks gradually suppress the male’s courtship behavior.

### **II.** Shock-induced courtship suppression persists for 7 days

We next wondered whether shock-induced courtship suppression is transient or persistent (Fig. 2a). To test this, we implemented the shock conditioning for 1 or 4 days, and tested the courtship behavior either 1 or 7 days afterward (Fig. 2b). We found that the suppression of orienting, singing, and mounting following 1-day conditioning consistently recovered 7 days afterward (Fig. 2c-e). In particular, shock-induced suppression of singing and mounting were detectable 1 day afterward (Fig. 2d, e) but undetectable 7 days afterward (Fig. 2d, e). We note that orienting tended to be suppressed 1 day after the conditioning but failed to reach the statistical significance (Fig. 2c), which is consistent with our previous observation (Fig. 1f). In stark contrast, following 4-day conditioning, courtship suppression persisted even 7 days afterward (Fig. 2c-e). This tendency was consistent across orienting, singing, and mounting, both in terms of the amount of time spent on, and the latency of initiating, each behavior (Fig. 2c-e). Together, our data suggests that repeated electric shocks persistently suppress male’s courtship behavior for at least 7 days.

**Fig. 2.**
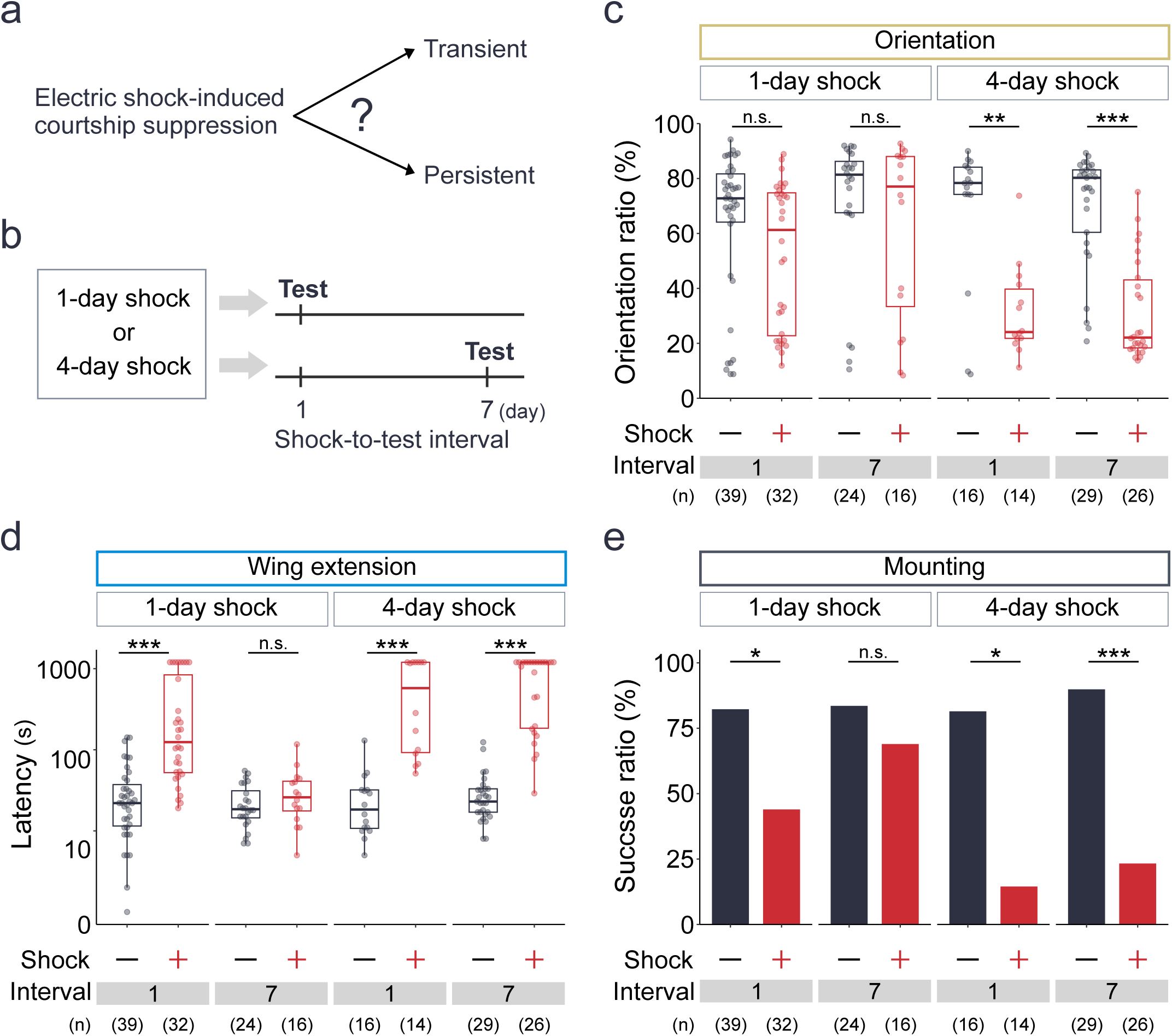
Shock-induced courtship suppression persists for 7 days. **a.** Hypotheses. **b.** Time course of the experiment. **c.** % time spent on orientation between the beginning of the recording and the onset of mounting (or the end of recording, if the male failed to mount). ***p < 0.001, **p < 0.01, n.s.: p > 0.05, two-way ANOVA followed by Tukey HSD. **d.** Latency of the initial wing extension in seconds. ***p < 0.001, n.s.: p > 0.05, two-way ANOVA followed by Tukey HSD. **e.** % flies mounted a female. ***p < 0.001, *p < 0.05, n.s.: p > 0.05, Fisher’s exact test with Bonferroni correction.

### III. Repeated shocks preferentially suppress P1 neurons in the courtship circuitry

We next sought to identify the mechanism underlying the courtship suppression. In Drosophila, the courtship circuitry from sensory first order neurons to the motor command neurons have already been mapped ^9,10^ (Fig. 3a), and advanced genetic toolbox allows for the recording and manipulation of each component of this circuitry. We thus set out to test whether repeated shocks might influence each of the neurons comprising this circuitry, using the recently developed activity monitoring technology^26^ (Fig. 3b). In this technology, a GFP-tagged CRTC (CREB-regulated transcriptional coactivator) is shuttled to the nucleus in response to the intracellular calcium increase, presumably due to neuronal activities. Therefore, by calculating the nucleus localization index (NLI) as (average GFP signals in the nucleus) – (average GFP signals in the cytosol) / (average GFP signals in the nucleus) + (average GFP signals in the cytosol). NLI thus ranges from -1 to 1, with the estimated neuronal activity being larger as NLI approaches 1. We expressed a GFP-tagged CRTC in each neuronal group comprising the courtship circuitry of male flies, and measured their activities 1 day after the 4-day conditioning, without prior interactions with a female. We found that, while activities of the first order (labeled by ppk23-GAL4), second order (labeled by AbdBLDN-GAL4), and song-triggering pIP10 neurons (labeled by pIP10 split-GAL4) failed to show significant differences between conditioned vs control flies, P1 neurons (labeled by P1 split-GAL4) showed a stark reduction in their activities following the shock conditioning (Fig. 3c; Supplementary Fig. 2a-d). This suppression was persistent, as P1 neurons’ activities remain suppressed even 7 days afterward (Fig. 3d; Supplementary Fig. 2e). Our data hints at the possibility that repeated shocks preferentially and persistently target the courtship-triggering P1 neurons to suppress courtship.

**Fig. 3.**
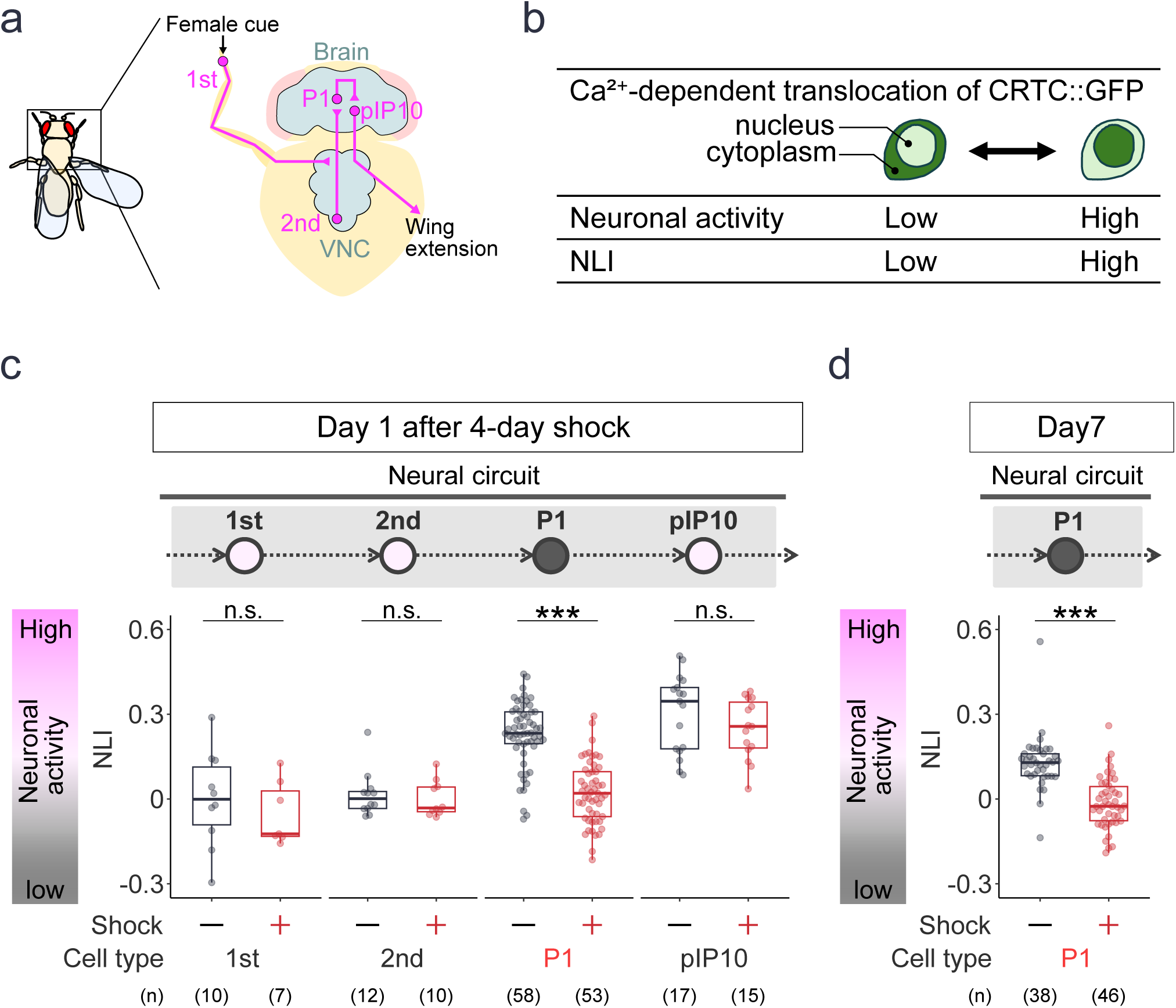
Repeated shocks preferentially suppress P1 neurons in the courtship circuitry. **a.** A schematic of the courtship circuitry. **b.** Activity monitoring technology used. A GFP-tagged CRTC (CREB-regulated transcriptional coactivator) is shuttled to the nucleus in response to the intracellular calcium increase, presumably due to neuronal activities. The degree of nucleus localization (nuclear localization index, NLI) thus reflects the neuronal activity level. **c.** NLI of the courtship circuitry 1 day after the conditioning. ***p < 0.001, n.s.: p > 0.05, two-sided Wilcoxon’s rank-sum test with Holm correction. **d.** NLI of the courtship circuitry 7 days after the conditioning. ***p < 0.001, n.s.: p > 0.05, Wilcoxon’s rank-sum test.

### **IV.** P1 neurons, but not pIP10 neurons, require stronger photoactivation to drive courtship after repeated shocks

Our data so far hints that repeated shocks preferentially inhibit the courtship-triggering P1 neurons to suppress courtship. We reasoned that, if this possibility is the case, then (1) optogenetic activation of P1 neurons would drive courtship even in conditioned flies, but (2) stronger light would be required for conditioned flies, compared to control flies, to drive courtship (Fig. 4a). To test this, we expressed in P1 neurons a light-gated cation channel CsChrimson, conditioned the flies for 4 days, and photoactivated their P1 neurons in the absence of female (Fig. 4b; Supplementary Fig. 3a). We found that photoactivation of P1 neurons induced singing (wing extension) in both control and conditioned flies, but that conditioned flies required stronger light intensity than unconditioned counterparts for singing (Fig. 4c; Supplementary Movie 3). More specifically, as we increased the light intensity, the fraction of control flies singing gradually increased, reaching a plateau at 1 mW/cm^2^ (Fig. 4c, black). In contrast, a plateau was reached for conditioned flies only at 4.3 mW/cm^2^ (Fig. 4c, red). This is consistent with the idea that the baseline activities in P1 neurons are suppressed following repeated shocks.

**Fig. 4.**
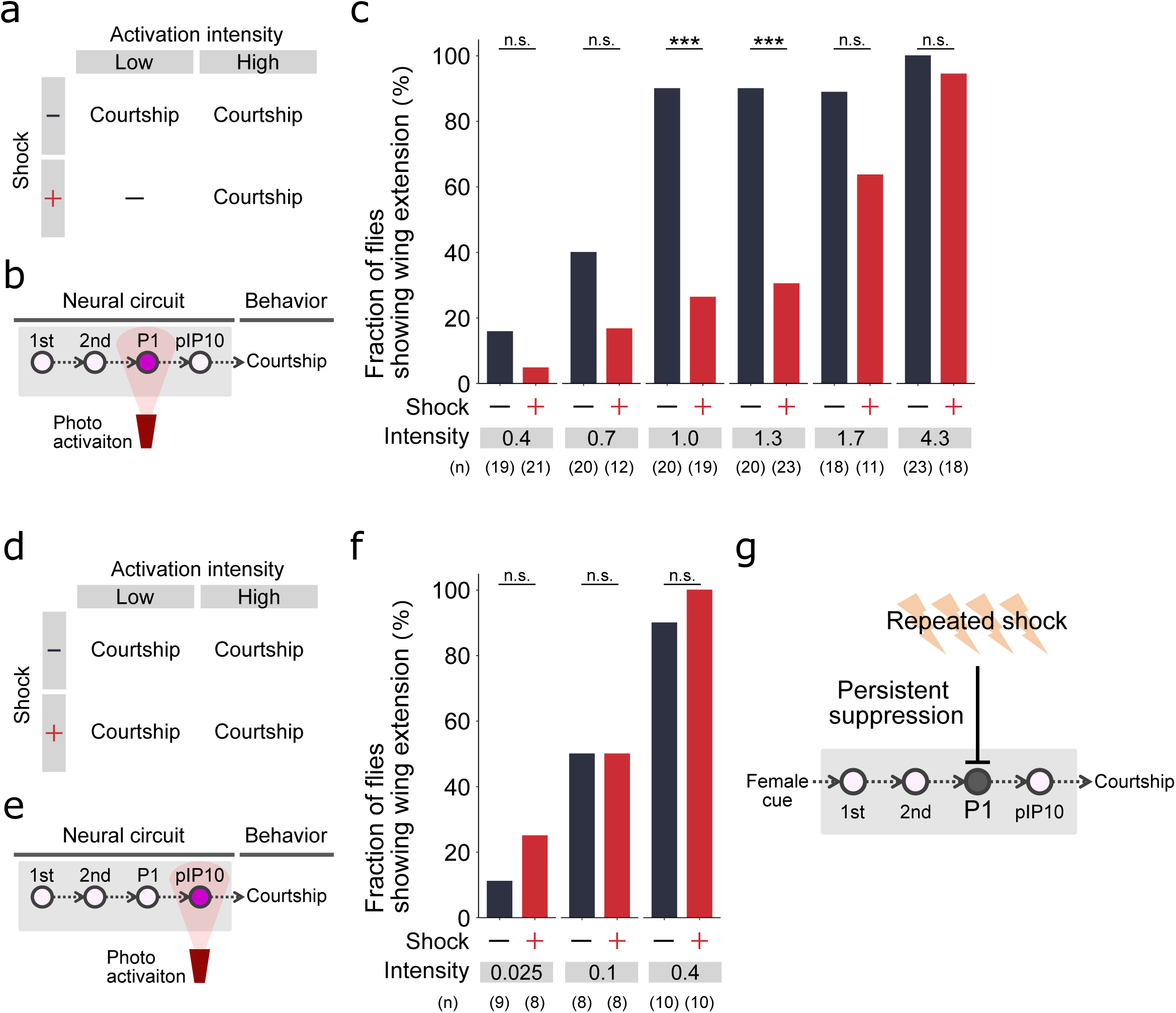
P1 neurons, but not pIP10 neurons, require stronger photoactivation to drive courtship after repeated shocks. **a.** Hypothesis. **b.** A schematic of the experiment. P1 neurons of a male fly were optogenetically activated without the presence of female, and the resultant courtship (wing extension) behavior was assessed. **c.** % flies showing wing extension. ***p < 0.001, n.s.: p > 0.05. Fisher’s exact test with Bonferroni correction. **d.** Hypothesis. **e.** A schematic of the experiment. pIP10 neurons of a male fly were optogenetically activated without the presence of female, and the resultant courtship (wing extension) behavior was assessed. **f.** % flies showing wing extension. ***p < 0.001, n.s.: p > 0.05. Fisher’s exact test with Bonferroni correction. **g.** Model.

Our model that shocks preferentially inhibit P1 neurons also predicts that photoactivation of neurons downstream of P1 neurons would induce courtship behavior in control and conditioned flies in the same manner (Fig. 4d). To test this prediction, we photoactivated the pIP10 neurons, post-synaptic to P1 neurons (Fig. 4e). We found that photoactivation of pIP10 neurons induced singing in both control and conditioned flies in a similar manner across different light intensities (Fig. 4f; Supplementary Movie 4), in agreement with our model.

Overall, the present study suggests that repeated shocks preferentially and persistently suppress P1 neurons to reduce courtship behavior (Fig. 4g). Our data illustrates how repeated stress preferentially targets a single node in the circuitry to persistently alter behavior.

## DISCUSSION

Here we have discovered that repeated electric shocks to male flies persistently suppress courtship behavior (Fig. 1, Fig. 2). We further found that repeated shocks preferentially suppress a group of courtship-triggering P1 neurons in the courtship circuitry (Fig. 3), which causally contributes to the courtship suppression (Fig. 4). Collectively, our data suggests a neuronal mechanism in which repeated stress persistently alters behavior by targeting a specific node of the behavior-driving circuitry.

### Electric shock as stressor in Drosophila

A stressor is often defined as external or internal stimuli that disturbs homeostasis and thus triggers compensatory biological responses^27^. In practice, it is operationally defined as any experimental manipulation that reliably produces a set of behavioral (escape, altered motivation), physiological (e.g., stress hormone elevation), and neural (e.g., amygdala/ parabrachial nucleus/ periaqueductal gray activation) responses, which can accumulate when stressors are applied repeatedly and/or for a long time ^2,3,27^. Our data shows that electric shocks elicit a set of flying, jumping, and running behavior (Supplementary Fig. 1b; Supplementary Movie 1), reminiscent of when the flies are exposed to a visual loom or noxious heat^23–25^, suggesting these behaviors indicate escape attempts. Furthermore, repeated exposure to electric shocks persistently suppressed courtship behavior, which became more pronounced as the function of repeat (Fig. 1). In mammals, too, stressors can persistently suppress courtship behaviors ^15–18^. While physiological or neural stress responses have not been established in Drosophila, our behavioral data are in line with the idea that electric shocks serve as stressor in Drosophila, as in the case of mammals^28^.

What is the sensory modality of electric shocks? In mammals, electric foot shock activates peripheral nociceptors, and is believed to be noxious^29,30^. It is thus plausible that electric shocks are noxious to adult flies as well. While nociception circuitry in adult Drosophila is still poorly defined, future identification of such circuitry would address this issue. Furthermore, it would also be interesting to see whether different kinds of stressors such as loom or noxious heat can also produce similar behavioral and physiological changes.

### Targeted and not distributed modulation of behavior-driving circuitry by repeated stress

In the present study, we took a wholistic approach and evaluated throughout the courtship circuitry with or without shock conditioning. This approach allowed us to pinpoint the site of modulation imposed by repeated stress, which turned out to be a single, courtship-triggering node of the circuitry, called P1 neurons.

Repeated stress to mammals can modulate an array of distinct brain regions, such as hippocampus, ventral tegmental area (VTA), amygdala, medial prefrontal cortex (mPFC), to name a few^27^. Manipulating neuronal activities in these regions can in some cases rescue behavioral alterations in stressed animals, suggesting that these physiological modulations are not merely correlational but causal to at least some stress-induced behavioral alterations. However, distributed nature of such modulations makes it challenging to experimentally assess each modulation’s contribution to behavior. In addition, there has been no instance where the circuitry underlying a certain behavior has been completely mapped from sensory input to motor output in mammals, leaving the question of whether the known instance of neuronal modulation can fully account for the behavioral alterations, or instead additional modulations elsewhere need to be considered to fully understand the physiological basis of stress-induced behavioral alterations. Our study presents a valuable model system where the site of modulation has been mapped in the complete circuitry.

Overall, we have discovered that repeated electric shocks persistently and preferentially suppress P1 neurons in the courtship circuitry. Because our finding provides the first experimental model of repeated stress in which the site of modulation has been mapped in the complete circuitry, we propose that our findings open up an exciting avenue to the interrogation of how repeated stress persistently alters behavior.

## ONLINE METHODS

### Fly stocks

The following strains were obtained from Bloomington Stock Center (Indiana University): ppk23-GAL4 (#93026), AbdBLDN-GAL4 (#55848), pIP10 split-GAL4 (#87691), UAS-CRTC (#99657), P1 split-GAL4 (P1-AD (#68837), P1-DBD (#69507).

### Electric shock conditioning

Flies were maintained on conventional cornmeal medium under a 9AM:9PM light/ dark cycle at 25°C and 60 +/- 5 % humidity. Flies were collected on the day of eclosion, and housed in a group of 9-10 for 3 days before testing. All experiments were carried out between 10 am and 1 pm at 25 +/- 0.8 C and 60 +/- 5 % humidity.

Our electric shock chamber was 60 x 10 x 3 mm acrylic chamber. An array of copper wires tiles the floor, so that “+” and “-“ wires alternate each other. Thus, the current flows through the fly only when the fly steps over both “+” and “-“ wires. The voltage was set to 60 V unless otherwise stated, to be consistent with previous reports (Hu et al., 2018). The inner walls and ceiling of the chamber were coated with silicone (crepolymate, KURE) to prevent flies from climbing.

A group of flies were transferred from a food-containing vial to our conditioning chamber by gentle aspiration. Immediately afterward, the copper wires tiling the floor were electrically charged. The currents flowing was monitored with amperemeter (Digital Tester, OHM). Flies’ behaviors during conditioning were videotaped from above at 15 fps. The conditioning was implemented for 3 hours. Immediately afterward, the flies were aspirated back to a food-containing vial and were allowed to recover until the conditioning resumed the next day.

### Courtship behavior analysis

An acrylic chamber (3 mm in height, 10 mm in diameter) was placed on a translucent white diffusion plate. A webcam was positioned above the chamber, and a white LED illuminator was placed below it. Male and female flies were gently transferred into the chamber using an aspirator. Behavioral recordings were acquired for 20 min after both the male and female were introduced into the chamber. Behavioral annotations related to courtship were performed using FlyTracker^31^. The criteria for each behavior were defined as follows:

Orientation: the female’s center of mass was within ± 60° of the male’s anteroposterior body axis, the distance between their centers of mass was ≤ 6 mm, and the fly’s speed was ≥ 0.1 mm/s.

Singing: the maximum wing extension angle was ≥ 60°. For flies that failed to show wing extension during the 20 min recording, latencies of the initial wing extension were treated as 20 min.

Mounting: annotated manually.

### Optogenetics

Flies expressing CsChrimson, a red-shifted channelrhodopsin variant, in P1 or pIP10 neurons were raised on food containing 500 µM all-trans retinal throughout the conditioning-to-test interval. A single male fly was gently transferred into the courtship assay chamber using an aspirator. After 20 min of acclimation, behavioral recordings were acquired for 5 min. Photoactivation was performed by delivering LED light (617nm, Thorlabs) to the whole courtship assay chamber at 1 Hz (duty cycle: 50%) for 5 min. The output for LED was measured with a power meter (S120C and PM120D, Thorlabs), and was adjusted to 0.025 – 4.3 mW/cm^2^, as indicated in Fig. 4. Light stimuli were triggered by python through an Arduino board connected to the LED driver. From the beginning of retinal feeding to the end of behavioral recording, the flies were kept in the dark throughout.

### Immunohistochemistry

The following antibodies were used: mouse mCherry (1:1000, Developmental Studies Hybridoma Bank), chicken anti-GFP (1:1000, Developmental Studies Hybridoma Bank), goat anti-chicken Alexa 488 (1:200, Molecular Probe #A11039), anti-mouse Alexa 555 (1:200, Molecular Probe #A21424). Whole brain immunohistochemistry was performed as described previously^32,33^. Briefly, brains were dissected out in 0.3 % PBST, and were fixed in 4 % paraformaldehyde/ PBS for 90 min at room temperature. Brains were then washed in 0.3 % PBST, for > 30 min, and were blocked in 5 % normal goat serum/ 0.3 % PBST for 30 min. Brains were incubated in the primary antibodies in 5% normal goat serum/ 0.3% PBST at 4°C for 1 day. Brains were then washed in 0.3 % PBST for > 30 min, and were blocked in 5 % normal goat serum/ 0.3 % PBST for 30 min. Brains were then incubated in the secondary antibodies in 5 % normal goat serum/0.3 % PBST at 4 for 1 day. Afterward, brains were washed in 0.3 % PBST for > 30 min, and were mounted on slide glass in Vectashield (Vector Laboratories). Images were taken with Leica TCS SP8 confocal microscope, and processed in ImageJ (NIH) software.

### Data analysis

Data analyses were performed using our custom-written Matlab, R, and python programs.

### Statistics

We did not predetermine the sample sizes. For one-sample tests shown in Fig. 3, non-parametric, Wilcoxon signed rank test was used; For Fig. 4c, f, Fisher’s exact test with Bonferroni correction was used; for Fig. 3c, d, Wilcoxon rank-sum test with Holm correction was used; for the other data, two-way ANOVA followed by Tukey HSD was used. All statistical tests were two-sided. Neither randomization nor blinding was performed for the group allocation during experiments or data analysis. All box plots are generated so that center line indicates median, box limits indicate upper and lower quartiles, and whiskers indicate 1.5x interquartile range.

## ACKNOWLEDGEMENTS

We thank T. Katsuki and T. Tachizaki (ThorLabs) for technical support in setting up the hardware; members of the Emoto Lab for advice and support. This study was supported by JST (JPMJSP2108) to Y.N., JSPS (KAKENHI 24K18157), JST (ACT-X JPMJAX242D), and Yamaguchi Educational and Scholarship Foundation to M.T. and AMED-CREST (JP21gm310010), Takeda Science Foundation, Torray Foundation, Naito Foundation, WPI-IRCN to K.E.

## AUTHOR CONTRIBUTIONS

Y.N., M.T., and K.E. designed the experiments. Y.N. and M.T. performed the experiments. Y.N. analyzed the data. Y.N., M.T., and K.E. interpreted the results and wrote the paper.

## FIGURE LEGENDS

**Supplementary Fig. 1 Repeated electric shocks and the courtship behavior assay**

**a.** An electric shock chamber. **b.** (Upper) An example time course of the shocks, occuring at random timings as the flies step over both “+” and “-“ wires. (Lower) An ethogram showing shock-induced flying, jumping, and running behaviors. **c.** A courtship assay chamber.

**Supplementary Fig. 2 CRTC results of the courtship circuitry**

**a.** Example CRTC images of 1^st^-order neurons (labeled by ppk23-GAL4) with or without shock conditioning. The images were acquired 1 day after the conditioning. **b.** Example CRTC images of 2^nd^-order neurons (labeled by AbdBLDN-GAL4) with or without shock conditioning. The images were acquired 1 day after the conditioning. **c.** Example CRTC images of P1 neurons (labeled by P1 split-GAL4) with or without shock conditioning. The images were acquired 1 day after the conditioning. **d.** Example CRTC images of pIP10 neurons (labeled by pIP10 split-GAL4) with or without shock conditioning. The images were acquired 1 day after the conditioning. **e.** Example CRTC images of P1 neurons (labeled by P1 split-GAL4) with or without shock conditioning. The images were acquired 7 days after the conditioning.

**Supplementary Fig. 3 Optogenetic induction of courtship behavior**

**a.** Red light was shined from above to the whole courtship assay chamber.

**Supplementary Movie 1**

An example batch of flies undergoing shock conditioning.

**Supplementary Movie 2**

Example pairs of flies engaging in the courtship behavior (left, a control male) or failing to engage in the courtship behavior (right, a conditioned male).

**Supplementary Movie 3**

An example, control male showing singing (wing extension) in response to photoactivation of P1 neurons.

**Supplementary Movie 4**

An example, control male showing singing (wing extension) in response to photoactivation of pIP10 neurons.

